# Inherited Human Sex Reversal due to Loss of a Water-Mediated Hydrogen Bond at a Conserved Protein-DNA Interface

**DOI:** 10.1101/2021.05.05.442830

**Authors:** Yen-Shan Chen, Joseph Racca, Dan Amir, Elisha Haas, Michael A. Weiss

**Affiliations:** From the Department of Biochemistry and Molecular Biology Indiana University, School of Medicine, Indianapolis, IN 46202; Faculty of Life Sciences, Bar Ilan University, Ramat Gan, Israel 52900

**Keywords:** sexual development, intersex, gonadogenesis, gene regulation, protein-DNA recognition

## Abstract

Male sex determination in mammals is initiated by SRY, a Y-encoded architectural transcription factor. The protein contains a high-mobility-group (HMG) box that mediates sequence-specific DNA bending. Mutations in SRY causing XY gonadal dysgenesis (Swyer syndrome) cluster in the box. Although such mutations usually arise *de novo* in spermatogenesis, some are inherited: male development occurs in one genetic background (the father) but not another (the sterile XY daughter). Here, we compare *de novo* and inherited mutations at an invariant Tyr adjoining the motif’s basic tail (consensus position 72; Y127C and Y127F in intact SRY). Crystal structures of homologous SOX-DNA complexes suggest that the wild-type side chain’s *para*-OH group anchors a water-mediated hydrogen bond to the DNA backbone. In an embryonic gonadal cell line, Y127C (*de novo*) led to accelerated proteasomal proteolysis and blocked transcriptional activity; activity remained low on rescue of expression by chemical proteasome inhibition. Y127F (inherited) preserved substantial transcriptional activity: 91(±11)% on SRY overexpression and 65(±17)% at physiological expression. Control studies indicated no change in protein lifetime or nuclear localization. Only subtle biophysical perturbations were observed *in vitro*. Although though inherited variant’s specific DNA affinity was only twofold lower than wild type, stopped-flow kinetic analysis revealed a sevenfold decrease in lifetime of the complex. Time-resolved fluorescence energy transfer (using a 15-base pair DNA site) demonstrated native mean DNA bending but with a slightly widened distribution of end-to-end DNA distances. Our findings highlight the contribution of a single water-mediated hydrogen bond to robustness of a genetic switch in human development.

---

The male phenotype in eutherian mammals is determined by *Sry* (1), an architectural transcription factor encoded by the Y chromosome (2). SRY contains a central high-mobility-group (HMG) box (Fig. 1), a conserved motif of specific DNA binding and DNA bending (3). This signature domain (*dark blue* in Fig. 1*A*) and its basic tail (*light blue*) are conserved among an extensive family of SRY-related transcription factors (SRY-related HMG box; SOX) broadly involved in development. The discovery of clinical mutations in SRY leading to 46, XY gonadal dysgenesis with female somatic phenotype (Swyer Syndrome) has provided evidence in support of SRY’s function in human testis determination (4,5). Such mutations cluster in the HMG box and usually occur *de novo* in paternal spermatogenesis (6,7).

**Figure 1.**
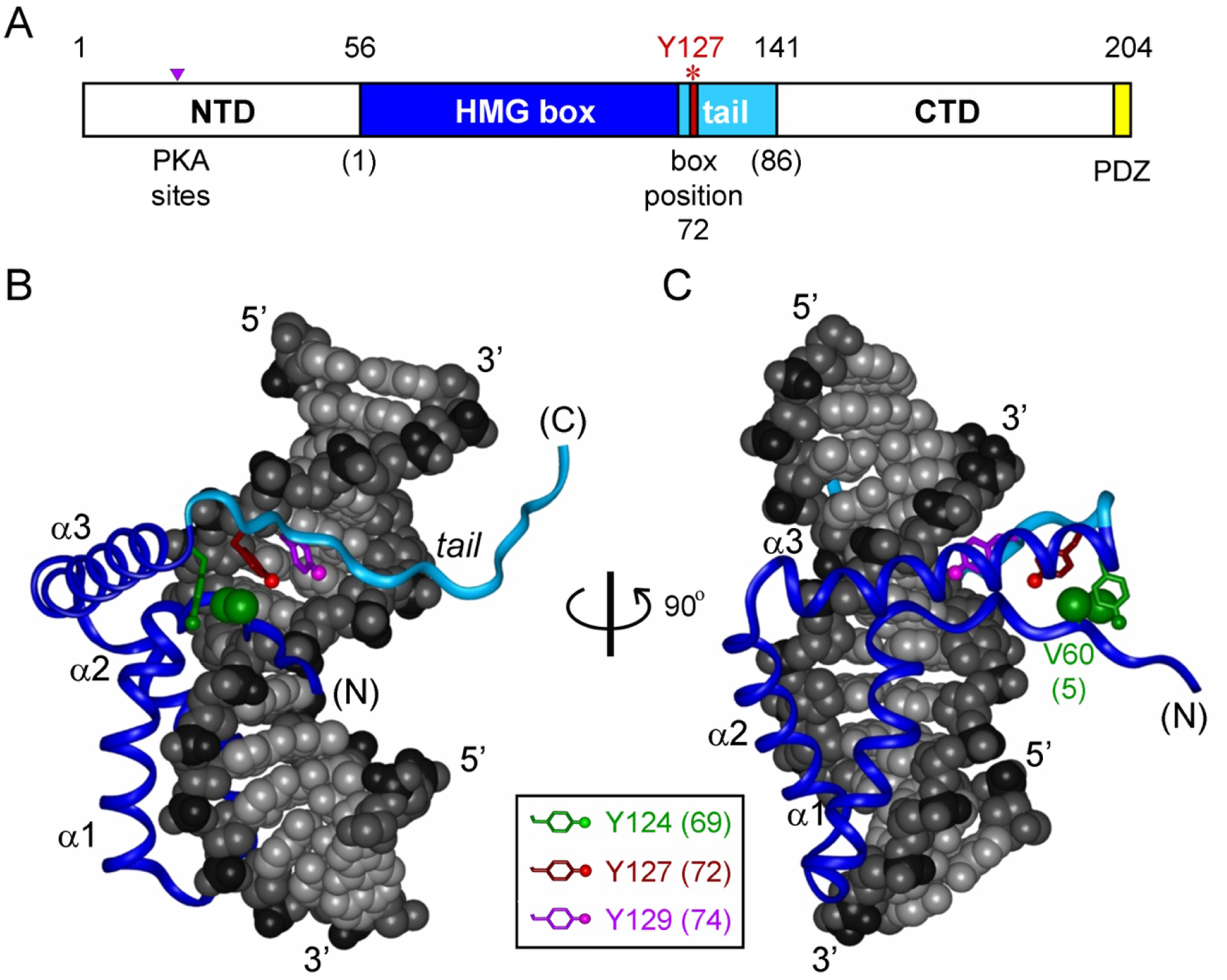
Domain organization of SRY and structure of DNA-bound complex. (*A*) Domains of human SRY and sites of tyrosine 127. The HMG box (*dark blue*) and its basic tail (*light blue*) comprise residues 56-141. The N-terminal segment domain (NTD) contains two potential phosphorylation sites modulating DNA binding (*purple triangle*; position 31-33, (66)) the extreme C-terminus (CTD) contains PDZ-binding motif (*yellow*, (67)), which is proposed to interact with SIP-1. Above the bar are indicated full-length numbers of SRY. Red asterisk highlights the tyrosine in full-length position 127 (consensus number 72). Below the bar are consensus HMG box residue numbers in parenthesis. *Not shown:* the bridge domain follows the HMG box and is proposed to mediate protein-protein interactions; HMG-box nuclear localization signals N-NLS and C-NLS. (*B*) Ribbon model of DNA-binding form of SRY HMG box; HMG box docks within minor groove of bent DNA site (space-filling representation). Highlighted in the ribbon are sites of basic tail (*light blue*). The domain contains major wing (consisting of α-helix 1, α-helix 2, the first two turns of α-helix 3, and connecting loops) and minor wing (the N-terminal β-strand, the remainder of α-helix 3, and C-terminal tail). In a specific complex an interface forms between α-helix 3 and the N-terminal β-strand to stabilize an L-shaped structure. The DNA atoms are dark gray (phosphodiester linkages), medium gray (deoxyribose moieties) and light gray (base pairs). The aromatic box is depicted with the side chains of V60, H120, Y124, and Y127. Tyrosine side chains are color-coded as defined in the inset box. Also shown is the position of V60 side chain (residue 5 in the HMG box), which in bound state packs in mini-core of minor wing. (*C*) The complex in panel *B* as viewed following a 90° rotation about the vertical axis. Although Y127 is not in contact with the DNA, its packing within the minor wing buttresses the DNA-binding surface. The consensus residue number is in parenthesis.

How SRY initiates testicular differentiation of the bipotential gonad has been investigated in mouse models (8). Murine *Sry* is expressed in the pre-Sertoli cells of the differentiating gonadal ridge; the protein binds to specific regulatory DNA sites within testis-specific enhancers (*TES and Enh13*) upstream of *Sox9*, an autosomal gene broadly conserved among vertebrate sex-determining pathways (9-11). Sox9 is itself a transcription factor, activating a male-specific gene-regulatory network (GRN) in the fetal gonadal ridge (10,11): this regulatory axis orchestrates programs of cell-cell communication, migration and differentiation leading to testis formation (12,13). Endocrine functions of the fetal testis then direct regression of the female anlägen (via secretion of glycohormone Muellerian Inhibiting Substance [MIS/AMH] (14,15)), and somatic virilization (via secretion of testosterone; (16)). Mutations at successive levels of this pathway are broadly associated with disorders of sexual development (DSD) (17,18).

Comparative studies of Swyer variants in SRY have demonstrated a range of biochemical and biophysical perturbations (19-23). Sporadic mutations in SRY are typically associated with marked impairment of specific DNA binding (24-27) whereas inherited mutations exhibit more subtle perturbations (28). The latter are by definition compatible with alternative developmental outcomes: testicular differentiation leading to virilization (fertile 46, XY father) or nascent ovarian differentiation leading to gonadal dysgenesis (sterile 46, XY daughter) (4,6,29-31). Inherited mutations in SRY thus provide “experiments of nature” that define mechanistic boundaries of genetic function. These boundaries may pertain to SRY’s specific DNA-binding properties (28) or to other cellular processes (32), such as nucleocytoplasmic shuttling (33), post-translational modification or proteasomal degradation (34).

The HMG box is an L-shaped domain containing an N-terminal β-strand (consensus residues 1-11), three α-helices (α_1_, α_2_ and α_3_) and a basic tail (Fig. 1*B* and *C*) (35). Packing of the β-strand against the C-terminal segment of α_3_ defines the minor wing in the HMG box, whereas α_1_, α_2_ and the N-terminal portion of α_3_ comprise the major wing. Unlike structure-specific homologs (such as the boxes of HMG-1 and HMG-D, well ordered in the absence of DNA (36,37), the minor wing of the SRY domain is partly disordered: its N-and C-terminal strands are unfolded, and the major wing is molten (38,39). Specific DNA binding within an expanded minor groove stabilizes the canonical L-shaped structure (40). The sequence-specific protein-DNA interface is remarkable for “wedge insertion” of nonpolar side chains in addition to canonical hydrogen bonds and salt bridges (41). Specific contacts in the SRY-DNA complex are conserved within the ancestral SOX family of eukaryotic transcription factors (42). The SOX-DNA interface is extended by a C-terminal basic tail (*blue* in Fig. 1*A*; (21,22)), which augments the kinetic stability of the bent complex (23).

The present study has focused on two clinical mutations at a conserved site at the junction between helix α_3_ and the basic tail: box position 72 (4,29,43,44). One mutation (Y127C in intact human SRY) occurred *de novo* (45) whereas the other (Y127F) was inherited (31). The wild-type (WT) side chain is of dual structural interest. On the one hand, one face of the aromatic side chain buttresses a DNA-binding surface of the protein (Fig. 1*B* and 2*A*; (21,28)) via a cluster of aromatic residues engaging V60 (box position 5): H120, Y124, and Y127 (respective box positions 65, 69 and 72). On the other hand, the *para*-OH group of Y127 faces the DNA phosphodiester backbone but too far for direct engagement (6.7 Å from the phenolic oxygen to the nearest non-bridging oxygen atom). Whereas Y127C would be expected to perturb the aromatic cluster—in turn leading to transmitted perturbations of its overlying DNA-binding surface—how and to what extent Y127F impairs SRY function has remained enigmatic.

A previous study has reported that Y172F blocks detectable binding of the SRY HMG box to a specific DNA site (31). If valid, this finding poses a paradox: whereas blocked DNA binding could rationalize the phenotype of the XY daughter, what of the fertile father? Indeed, inheritance of this variant allele implies that Y127F SRY is compatible with alternative developmental outcomes: fertile father or sterile daughter. To resolve this seeming paradox, we undertook comparative cell-based and biophysical studies of Y127C and Y127F SRY. Whereas the *de novo* mutation markedly impairs SRY function, effects of Y127F are modest: substantial gene-regulatory function is retained in accordance with its inheritance. Only subtle biophysical differences were observed, relative to the WT complex, by ^1^H-NMR and time-resolved fluorescence energy resonance transfer (tr-FRET). Molecular modeling based on homologous SOX-DNA crystal structures (46,47) suggests that the *para*-OH group of the conserved Tyr anchors a water molecule at the protein-DNA interface. We envision that loss of this single water-mediated hydrogen bond “unlocks” the bent protein-DNA complex and attenuates the male switch to the edge of ambiguity.

## RESULTS

An immortalized embryonic pre-Sertoli cell line (rodent XY cell line CH34 (20,48)) was employed in transient-transfection assays to probe SRY-directed transcriptional activation of SRY’s principal target gene, *Sox9*, in its native chromosomal context (28). Employing quantitative reverse-transcriptase PCR (qPCR), this assay measured time-dependent accumulation of mRNAs encoded by downstream genes in the SRY-*Sox9* GRN (49) following transient transfection of WT or mutant *SRY* constructs. Level of expression of the transfected SRYs were made high (overexpression; >10^6^ protein molecules per cell) or low (physiological expression; 10^3^-10^4^ molecules per cell) by an empty-plasmid dilution protocol (33).

### Transcriptional activities and stabilities of SRY variants

CH34 cells were transiently transfected with WT or variant SRY-encoded plasmids. Transfection efficiency was 42(±1)% as monitored by control co-transfection of a plasmid encoding green fluorescent protein (GFP). Transient transfection of WT SRY under standard conditions (1 μg plasmid DNA/well) augmented expression of *Sox9* by eightfold relative to the empty vector (*black bar* in Fig. 2*A*). Extent of transcriptional activation decreased on successive dilution of the SRY-encoded plasmid by the empty vector (a protocol that maintains constant total transfected DNA) to a final dilution of 0.02 μg SRY plasmid and 0.98 μg empty vector per well (50-fold dilution; *white bar* in Fig. 2*A*). Such fiftyfold dilution, typically yielding *ca*. 10^4^ protein molecules per cell in the absence of changes in rate of proteasomal degradation, provided a control for potential artifacts well known to be associated with transcription-factor overexpression.

**Figure 2.**
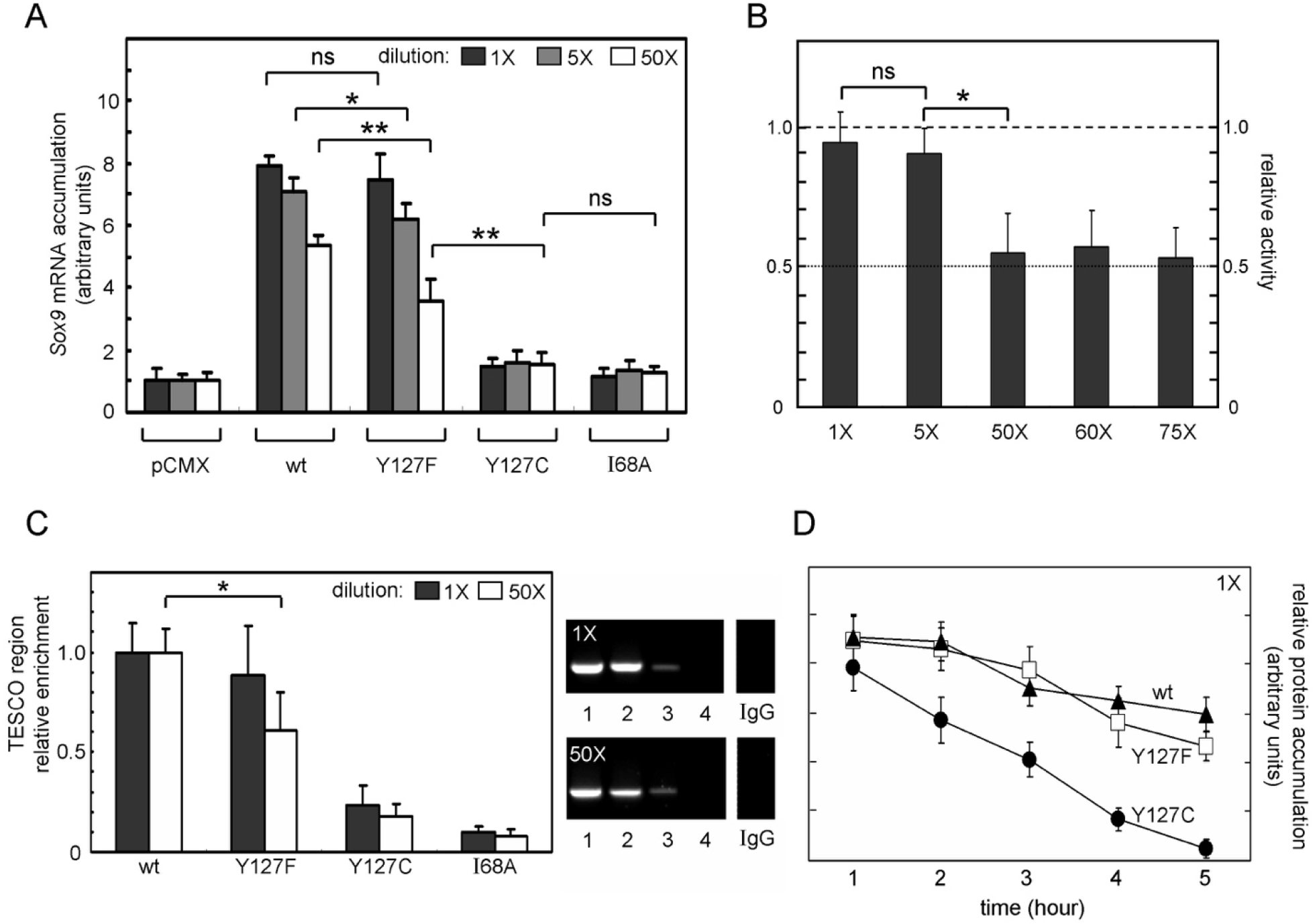
Assays of transcriptional activity. (*A*) Dependence of *Sox9* gene activation (monitored by *Sox9* mRNA accumulation) on dose of transfected plasmid SRY DNA: 1 μg (*black*), 0.2 μg (*dark gray*), and 0.02 μg (*white*). In each case total transfected DNA was the same. Horizontal brackets designate statistical comparisons: (* or **) Wilcox *p*-values <0.05 or 0.01 whereas “ns” indicates *p*-values > 0.05. (*B*) Relative activity (activity of Y127F comparing to that of WT SRY) in panel *B* under 1X, 5X, 50X, 60X and 75X-dilution of transfected Y127F SRY plasmid conditions. Statistical comparisons are defined in panel *A*. (*C*) ChIP assays probing SRY occupancy of target sites within *Sox9* testis-specific enhancer core element (TESCO). *Left*: Quantitative analysis of relative enrichment of SRY variant occupancies in selected dilution conditions (1X (no dilution) and 50X dilution in transfected SRY-encoded plasmids). Mutation which blocks specific DNA binding (68), I68A, serves as negative control. Statistical comparisons: (*) Wilcox *p*-values <0.05. *Right*: Gel showing results of ChIP of variants. Clinical mutations Y127C exhibit impaired TESCO region occupancies in all conditions but that of Y127F are only significant under 50X dilution of transfected SRY-encoded plasmid. (*D*) Cycloheximide assay. Mutation Y127F does not affect degradation rate significantly; proteolysis is enhanced by Y127C. This assay is under no-dilution plasmid condition (1X).

Initial investigation of the two variant SRYs (Y127F and Y127C) demonstrated that (i) Y127C markedly impaired transcriptional activity at each dilution whereas (ii) Y127F SRY exhibited native activity when overexpressed but partial loss of function on dilution (Fig. 2*B*). Remarkably, at the highest dilution (50-fold; white bars in Fig. 2*A*) Y127F was associated with approximately twofold loss of *Sox9* activation relative to WT SRY (Fig. 2*B*; 50X, 60X, and 75X). To investigate SRY-enhancer interactions, we employed chromatin immunoprecipitation (ChIP) to probe the testis-specific enhancers of *Sox9*. Initial studies focused on TESCO sites (10). Control studies of inactive I68A SRY (with Ala-shaved “cantilever” (50)) provided a baseline for complete loss of specific DNA binding (*right* in Fig. 2*C*). Whereas on overexpression of the proteins (*i*.*e*., without plasmid dilution) enhancer occupancies by WT and Y127F SRY were statistically indistinguishable (*black* bar in Fig. 2*C*), at a physiologic expression level the variant’s ChIP band intensities were reduced to *ca*. half of WT (Fig. 2*C*; *white* bar). By contrast, Y127C led to a marked decreased in enhancer occupancy under either transfection condition. Similar results were obtained in studies of a critical far-upstream enhancer region (*Enh13*; (11)) as shown in Supplemental Figure S1.

Although *Sox9* is the principal direct target of SRY (Fig. 3*A*, *boxed in red*), additional qPCR assays were undertaken to investigate a downstream GRN (activated by the Sox9 protein sequential to or in synergy with SRY); the studies were designed based on prior *in situ* transcriptional profiling of the XY gonadal ridge (51). Whereas transient expression of SRY did not affect mRNA accumulation of non-sex-related *Sox* genes and unrelated housekeeping genes (Fig. 3*B*), specific up-regulation of *Sox9, prostaglandin D2 synthetase* (*Ptgd2*) and *fibroblast growth factor 9* (*Fgf9*) was observed in accord with their known roles in testicular differentiation (Fig. 3; (52)). No such changes in mRNA accumulation were observed on transient transfection of an empty plasmid or a control plasmid expressing a stable SRY variant devoid of specific DNA-binding activity (I68A as above; (50)).

**Figure 3.**
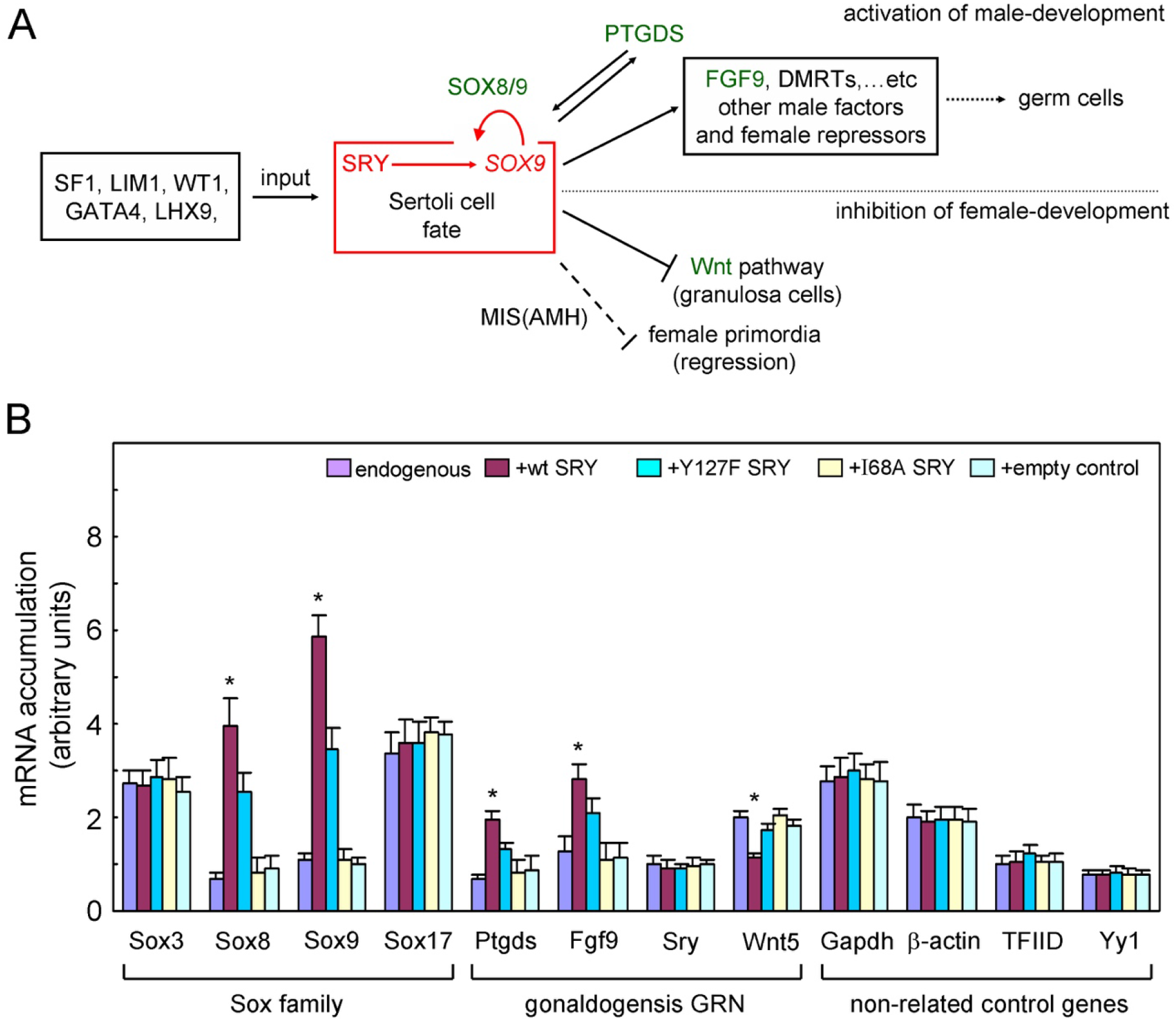
SRY and the male developmental gene-regulatory network (GRN). (*A*) The central regulatory axis of SRY→*SOX9* for the Sertoli cell differentiation (*red* box) with genetic input factors (box at *left*) and outputs (box at *right*) to activate a male-specific GRN (*upper*). This process also inhibits granulosa-cell fate (solid ⊥; Wnt pathway), and Müllerian regression (dashed ⊥; by MIS(AMH)). Red curved arrow indicates SOX8/9 feedback regulation to maintain *SOX9* expression once the SRY→*SOX9* regulation initiated. A*bbreviations:* FGF9, fibroblast growth factor 9; GATA4, GATA binding protein 4; LHX9, LIM homeobox 9; LIM1, homeobox protein Lhx 1; MIS (AMH), Müllerian Inhibiting Substance (Anti-Müllerian Hormone); PTGDS, Prostaglandin D2 synthase; WT1, Wilm’s tumor 1; Wnt, wingless-type. Color green highlights the selected gene for further investigation at panel *B*. (*B*) Selected gene expression patterns activated by various SRY variants in CH34 cell line. RT-Q-rt-PCR was employed to probe mRNA abundances of *Sox* family members (*left*), selected genes in gonadogenesis GRN (*middle*), and sex-unrelated housekeeping genes (*right*). qPCR was analyzed following expressions of SRY variant plasmids or control vectors including empty plasmid and a stable but inactive SRY variant (I68A). *Left*: Fold mRNA accumulation of *Sox* family genes, including *Sox8* and *Sox9* previously implicated in the program of SRY-mediated GRN. *Middle*: Fold mRNA accumulation of non-*Sox* sex-related factors. Results show that CH34 cells express low endogenous levels of *Sry, Fgf9, Ptgds*, and *Wnt5* gene expression. *Right*: Fold mRNA accumulation of sex-unrelated genes; these genes were not affected. Statistical analyses: *Wilcoxon test, *p*<0.05, indicating the significant comparison between WT and Y127F SRY transfections.

To characterize potential mechanisms, we further probed the proteolytic stabilities and subcellular localization of the WT and mutant proteins. Cellular turnover of the expressed epitope-tagged SRY proteins (transfected without dilution) was evaluated following treatment of cycloheximide, a chemical inhibitor of translation (Fig. 2*D*). Comparison of anti-hemagglutinin (Hal the epitope tag) Western-blot intensities demonstrated that the Y127C variant is more susceptible to degradation than is WT or Y127F SRY (Fig. 2*D*). Such differential degradation could be circumvented through addition of chemical proteasome-inhibitor MG132 to equalize expression levels (not shown).

### Nucleocytoplasmic localization of SRY variants

Subcellular localization was investigated using immunofluorescence microscopy (Fig. 4). To achieve statistical significance, 700 cells were counted in each case (performed in triplicate by blinded coworkers). Representative images are shown in Figure 4*A* (SRY; lower panel) in relation to corresponding nuclear staining of the same cells with 4’,6-diamidino-2-phenylindole (DAPI; upper panel). WT SRY and the Y127F variant exhibited similar subcellular distributions (Fig. 4*B*). Whereas Y127C without any additional treatment exhibited dramatic loss in both nuclear and pancellular staining (*gray* and *white* bars in Fig. 4*B*), rescue of its expression with native nuclear localization was achieved by treatment of proteasome inhibitor MG132 for 24 h post-transfection in serum-rich medium (“+MG132” at *right* in Fig. 4*B*). Evidently, these substitutions near a conserved nuclear localization signal (NLS; Fig. 4*C*) does not impair its import efficiency. Notably, addition of MG132 does not rescue the transcription potency of Y127C SRY (Fig. 4*D*). This result indicates that near-complete loss of Y127C SRY function is *intrinsic to its perturbed structure* and is not secondary to accelerated degradation of an otherwise active variant (34).

**Figure 4.**
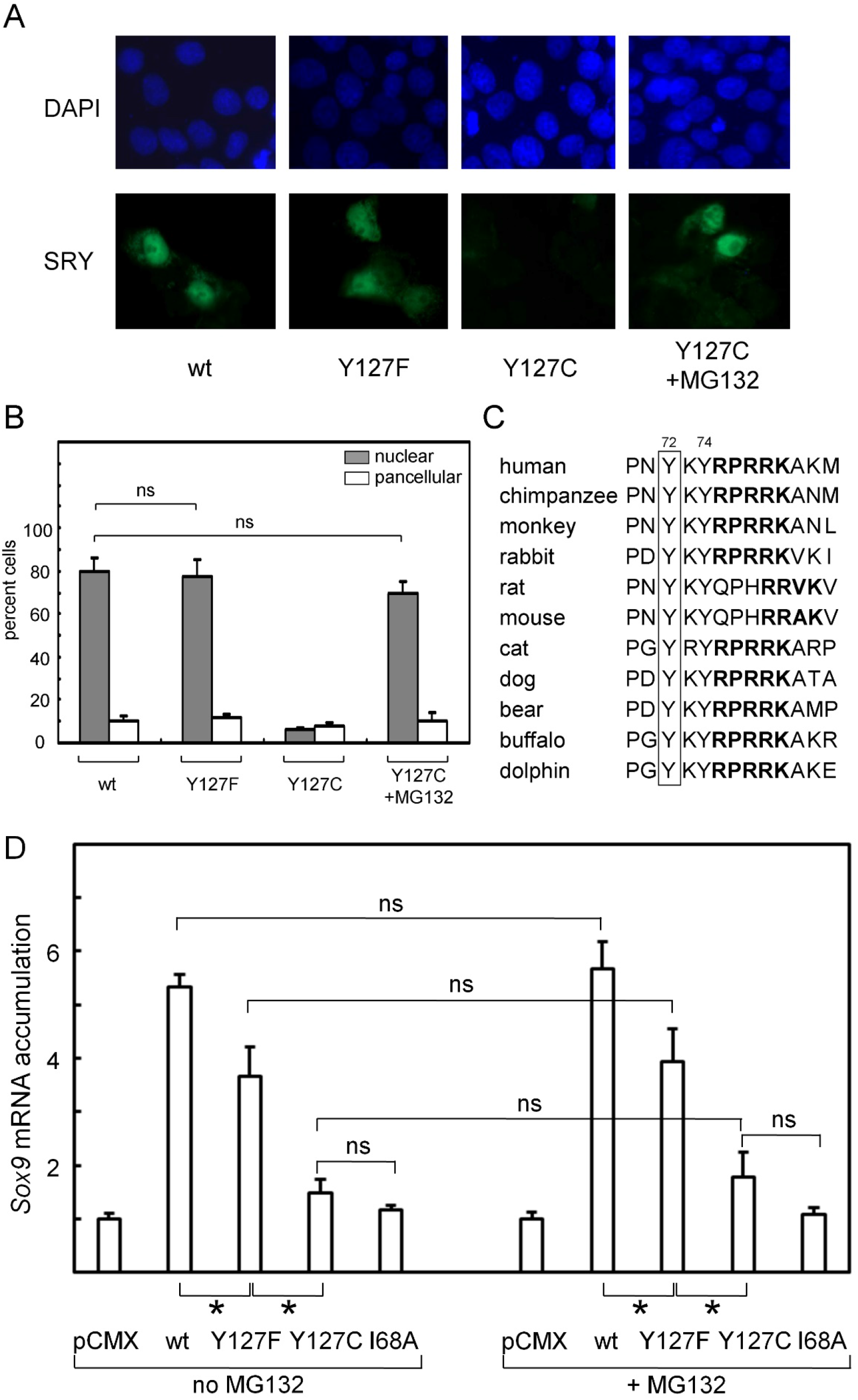
Assays of nuclear localization. (*A*) Subcellular localization of epitope-tagged SRY constructs as analyzed by immunostaining: DAPI nuclear staining (upper row *blue*) and SRY (lower row *green*). WT and Y127F SRY accumulate mainly in nucleus but distribute pancellularly; mutation Y127C SRY exhibits barely accumulation and could be rescued by MG132 treatment. (*B*) Histograms describing nuclear localization (*gray*) and pancellular pattern (*white*). Gray and white bars do not add to 100 due to occasional GFP-positive cells lacking SRY expression. SRY plasmid was at highest dose described in Figure 2*A*. Horizontal brackets indicate statistical comparisons as described above. (*C*) Sequence conservation of the minor wing of the SRY HMG boxes from various species. Tyr is invariant at position 72 of the HMG box among various mammalian *Sry* alleles (*boxed*). Residue numbers in an HMG-box consensus are provided above, and bold fonts highlight the C-terminal nuclear localization signal (NLS) in various species SRY. (*D*) CH34 cell line as treated with chemical proteasome inhibitor MG132. Histograms describing the *Sox9* gene expression in absence (left) or presence (right) of MG132. Data indicate that in no case did MG132 alter relative extent of *Sox9* expression; *i*.*e*., the impaired transcriptional activity of Y127C SRY could not be rescued by prevention of its proteasomal degradation. Statistical analyses: “ns” indicates no significant difference, asterisks at bottom indicate *p*<0.05; p-values (from left to right) are 0.036, 0.027, 0.046, 0.033 as calculated between WT and Y127F/C SRY transfections.

### Yeast-1-Hybrid Screening

Specific DNA binding by the Y127C and Y127F variant SRY domains was studied in a yeast-one-hybrid (Y1H) model system. An integrated reporter cassette (constructed to express β-galactosidase under the control of triplicate consensus SRY-binding sites) was employed. Expression of the reporter was regulated by a plasmid-encoded fusion protein containing the transcriptional activation domain of Gal4-linked to the WT HMG box or a variant. Whereas previous gel-mobility-shift assays (GMSA) of Y72F- and Y72C SRY (31,45) had shown no detectable specific DNA binding in either case, in the Y1H system the Y72F fusion protein gave rise to native transcriptional activity (Fig. 5*A*; *red*). By contrast, the Y72C fusion protein exhibited a marked loss of transcriptional activity (Fig. 5*A*; *gray*). These findings strongly suggested that specific DNA-binding activity was markedly impaired by Y72C but not by Y72F; due to overexpression of the fusion proteins in the yeast strain, small decrements in the specific DNA-binding affinity of the latter variant could not be excluded by these results.

**Figure 5.**
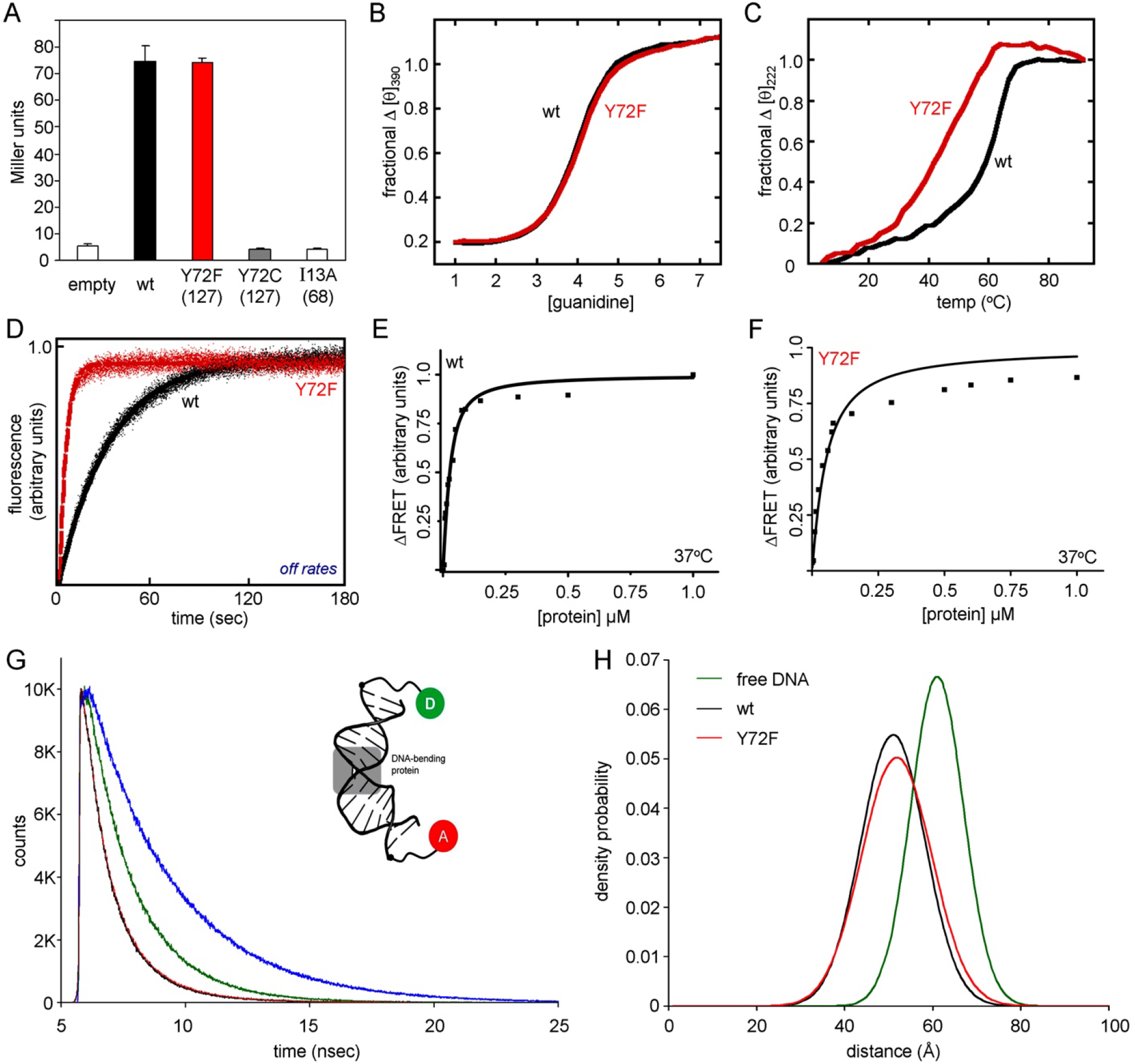
Biophysical and biochemical studies. (*A*) Yeast one hybrid assay exhibiting activation of an HMG box-dependent integrated reporter gene (expressing β-galactosidase) is similar between the WT and inherited Y72F mutant domain (full length SRY numbering in parentheses). *De novo* mutant, Y72C, exhibits no activation compared an empty vector or a variant HMG box (I13A) void of DNA binding. (*B* and *C*) Stability of the free (B) protein domains is similar, whereas the mutant protein-DNA complex (*C*) less stable relative to WT. (*D*) Kinetic stability studies of WT and Y72F mutant indicate the Y72F mutant domains has accelerated dissociation from its DNA site. Kinetic trace is representative measured at 15 °C, see Table 2 for all measurements. (*E* and *F*) Specific DNA binding at physiological temperature for the WT (*E*) and the mutant domain (*F*) as determined by FRET. The Y72F mutant has attenuated specific DNA binding. (*G* and *H*) Time resolved-FRET used to monitor the decay of the donor fluorophore on a nanosecond time scale, a schematic of the fluorescently-labeled DNA substrate used shown in (*G*) insert where the donor (5’-6FAM) is green and acceptor (5’-6TAMRA) is in red. The decay curves overlap for the WT and mutant, black and red lines respectively, compared to controls (single-labeled donor only in blue and free double-labeled DNA substrate in green). (*H*) Dynamic stabilities of the mutant and WT complex are similar, slight increase in peak width was measured for Y72F; Gaussian distribution of the end-to-end distances in the free (green) and bound DNA (red and black).

### Fluorescence studies of specific DNA binding and DNA bending

In the absence of DNA, the stabilities of the WT and mutant HMG boxes are similar as probed by chemical denaturation studies (Fig. 5*B* and Table 1). The stability of the protein-DNA complex was further investigated by circular dichroism (CD; Fig. 5*C*). Ultraviolet CD spectra of equimolar protein-DNA complexes were acquired at successive temperatures (in the range 4-91 °C). Midpoint melting temperatures (T_m_) were derived from analysis of ellipticity at helix-sensitive wavelength 222 nm. The apparent thermal stability of the Y127F complex (red in Fig. 5*C*; T_m_ 50 °C) was lower than that of the WT complex (black; *ca*. 63 °C) as summarized in Table 1. This decrement may in part reflect reduced protein-DNA affinity (below). Y72F thus affects the stability of the protein–DNA complex but not the free domain.

**Table 1.** Stability and DNA binding parameters

| sample | $\Delta G$<br>(kcal/mol) | $C_{mid}^a$<br>(M) | $T_m$ (°C)<br>complex | $K_D$ (nM)<br>15 °C | $K_D$ (nM)<br>37 °C |
| --- | --- | --- | --- | --- | --- |
| WT | 4.8 ± 0.12 | 3.1 ± 0.10 | 62.7 | 14.2 ± 2 | 14.5 ± 2 |
| Y72F | 4.4 ± 0.10 | 3.0 ± 0.10 | 50.2 | 21.7 ± 3 | 40.4 ± 4 |

<sup>a</sup> $C_{mid}$ represents that concentration of guanidine where the domain is half unfolded.

Specific protein-DNA dissociation constants (K_D_) were determined by an equilibrium method exploiting fluorescence resonance energy transfer (FRET) as previously described (28). The 15-base pair (bp) donor/acceptor-labeled DNA duplex containing the SRY consensus target site (5’-ATTGTT and complement) was held a constant concentration (25 nM), and energy transfer from the donor fluorophore (fluorescein) to the acceptor (tetramethylrhodamine; TAMRA) was measured at increasing protein concentrations; the assays were performed at both 15 and 37 °C. As shown in Table 1, the K_D_ for the WT HMG box does not change significantly between 15 and 37 °C (ca. 14-15 nM). The dissociation constant of the Y127F domain was weakened by 1.5-fold at 15 °C (K_D_ 22 nM) and 2.8-fold at 37 °C (K_D_ 40 nM; Fig. 5*E, F* and Table 1).

The same FRET-labeled DNA duplex enabled measurement of off-rate constants (*k*_*off*_) using a stopped flow kinetic FRET assay (23). An equimolar complex of protein bound to a fluorescently labeled DNA site was loaded into a syringe; a second syringe contained the corresponding unlabeled DNA duplex at 20-fold excess. Upon rapid mixing, the recovery of donor fluorophore was monitored as protein redistributes from the FRET-labeled DNA site to the excess unmodified DNA site in the mixing chamber. Representative data and its analysis are shown in Figure 5*D*. The results demonstrate that the Y127F domain more rapidly dissociates from the specific DNA site than does the WT domain. This assay was performed at temperatures 6, 15, 25 and 37 °C (Table 2). At each temperature the variant domain’s *off*-rate was greater than the WT domain’s *off*-rate. Because K_D_ = *k*_*off*_/*k*_*on*_, these data imply that the *on*-rate of the variant domain is also greater than that of the WT domain (although to a less extent).

**Table 2.**
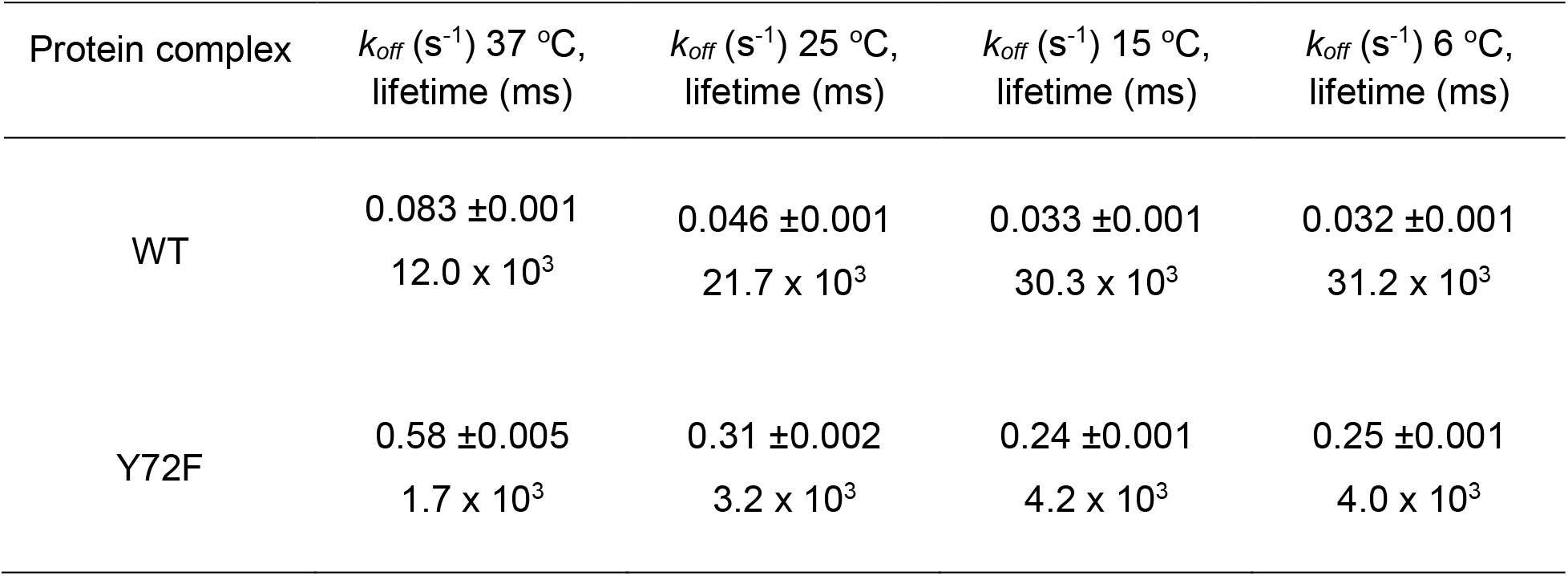
Off rate (*k*_*off*_) measurements by stopped flow FRET.

| Protein complex | $k_{off}$ (s<sup>-1</sup>) 37 °C,<br>lifetime (ms) | $k_{off}$ (s<sup>-1</sup>) 25 °C,<br>lifetime (ms) | $k_{off}$ (s<sup>-1</sup>) 15 °C,<br>lifetime (ms) | $k_{off}$ (s<sup>-1</sup>) 6 °C,<br>lifetime (ms) |
| --- | --- | --- | --- | --- |
| WT | 0.083 ±0.001<br>12.0 x 10<sup>3</sup> | 0.046 ±0.001<br>21.7 x 10<sup>3</sup> | 0.033 ±0.001<br>30.3 x 10<sup>3</sup> | 0.032 ±0.001<br>31.2 x 10<sup>3</sup> |
| Y72F | 0.58 ±0.005<br>1.7 x 10<sup>3</sup> | 0.31 ±0.002<br>3.2 x 10<sup>3</sup> | 0.24 ±0.001<br>4.2 x 10<sup>3</sup> | 0.25 ±0.001<br>4.0 x 10<sup>3</sup> |

Protein-directed DNA bending was investigated by time-resolved FRET (tr-FRET). In tr-FRET fluorescence decay of the donor was measured on a subnanosecond time scale; measurements using single-labeled DNA sites (donor only), double-labeled DNA sites (donor and acceptor) enable calculation of end-to-end distance distributions. The decay rates are similar for the WT and Y127F domain-DNA complexes (Fig. 5*G*, black and red, respectively); controls are shown in green (donor only) and blue (double-labeled dsDNA free). Global fitting of the decay curves yielded the mean end-to-end distances and distance distribution (Fig. 5*H* and Table 3). Although the inferred means were similar for the Y72F (red in Fig. 5*H*) and WT (black) complexes (in accordance with steady-state FRET; above), the mutation is associated with a small increase in the width of the distance distribution. Such broadening implies that a greater range of DNA bend angles and/or a greater range of DNA unwinding occurs in the variant complex relative to WT.

**Table 3.** Time-resolved FRET analysis of end-to-end DNA distances^a^

| sample | peak (Å) | mean (Å) | fwhm<sup>b</sup> (Å) | global $\chi^2$ |
| --- | --- | --- | --- | --- |
| DNA only | 60.7 (60.4 – 61.1) | 60.7 (60.4 – 61.1) | 18.2 (17.9 – 20.5) | 1.05 |
| WT | 51.1 (50.3 – 51.8) | 51.0 (50.3 – 51.7) | 28.9 (27.9 – 30.0) | 1.10 |
| Y72F | 51.6 (50.3 – 52.9) | 51.7 (50.6 – 52.8) | 32.9 (30.0 – 29.7) | 1.12 |

<sup>a</sup>Measurements were taken at 4 °C. The values in parentheses represent possible errors in parameters calculated. Methods are described in detail in Ref (64).
<sup>b</sup>Relative values of *fwhm* (full width at half max) provide an estimate of the precision of DNA bending but absolute values contain contributions from the flexibility of the linkers joining the fluorophores to the 5'-ends of the DNA duplex as observed in the free DNA (whose length is >tenfold shorter than the persistence length of B-DNA; ~50 nm or 150 bp).

## DISCUSSION

The interaction between the SRY HMG box and its specific DNA target site (core consensus sequence 5’-ATTGTT and complement) is remarkable for bidirectional induced fit: (i) protein-directed DNA bending in association with a widened and underwound minor groove; and (ii) DNA-directed folding of the minor wing of the HMG box in association with stabilization of the major wing and docking of the basic tail (21,22). In the free box, the N-terminal segment detaches from helix α_3_; the tail is disordered. The minor wing’s mini-core stabilizes—and is stabilized by—an overlying protein-DNA interface. Critical to such bidirectional induced fit is insertion of the isopropryl side chain of Val60 (box position 5) into a cluster of aromatic side chains: His120, Tyr124 and Tyr127 (respective box positions 65, 69 and 72; Fig. 6*A*). Paired proline residues C-terminal to α_3_ orient Tyr74 and successive basic residues to further stabilize the bent DNA complex. Mutations in these structural elements (variously perturbing mini-core packing, tail orientation or direct side chain/DNA contacts) result in gonadal dysgenesis and somatic sex reversal (28,31,53).

**Figure 6.**
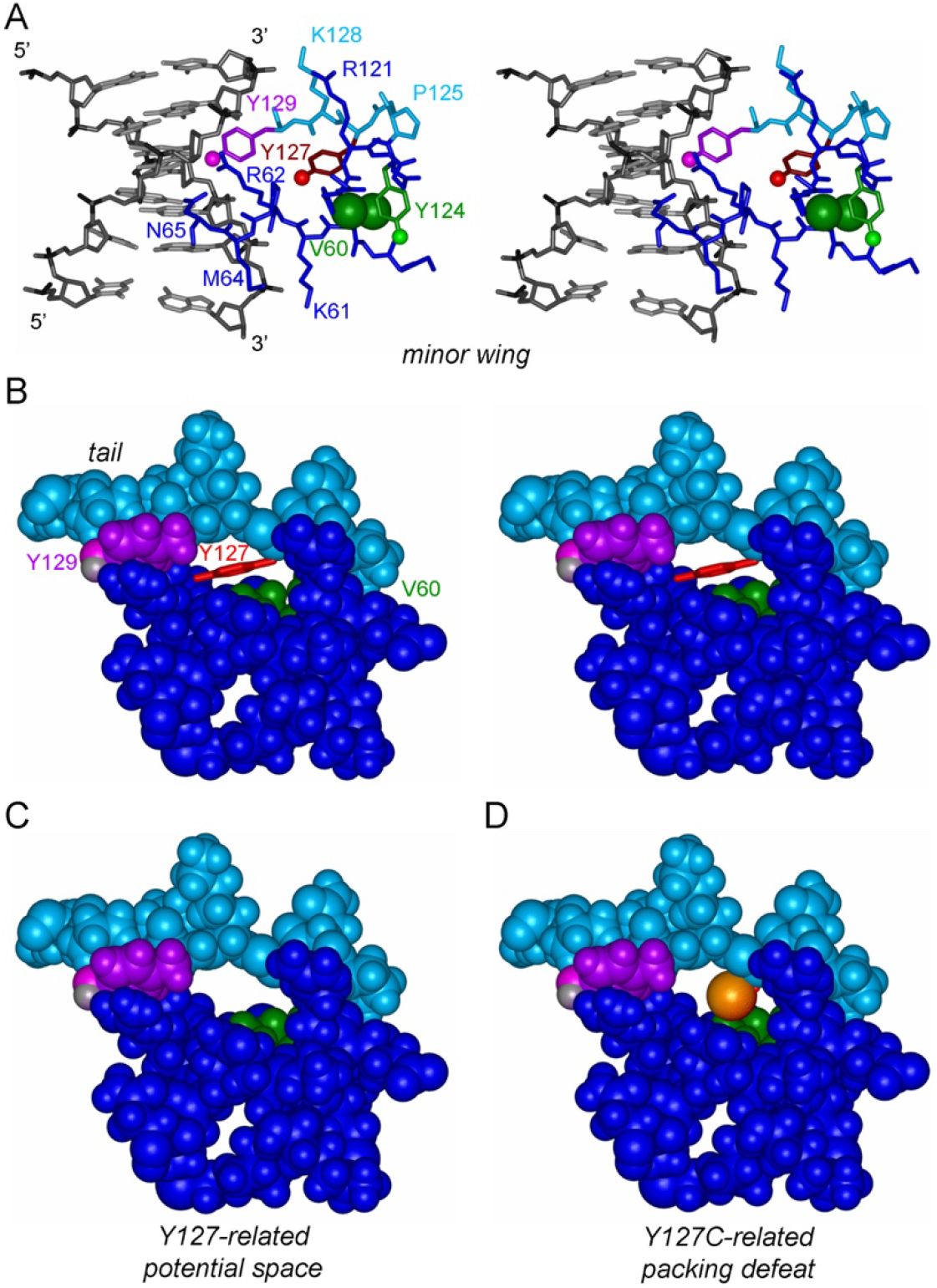
Overview of SRY minor wing. (*A*) Stereo pair of minor wing region in Figure 1*B* showing side-chain packing within the minor wing and protein-DNA contacts. Side chains and related color codes are same as definition in Figure 1*B*. (*B*) Stereo pair of space-filling model representation of the minor wing in the protein-DNA complex (DNA not shown). The side chain of Y127 is shown (red sticks) relative to the space-filling surface of the minor wing. The hydrogen atom in Y129 is highlighted with gray. (*C*) Space-filling representation showing a proposed model of minor wing in absence of Y127 side chain, and replaced with cysteine (*D*).

The present study has focused on two clinical mutations at residue 127 of human SRY (consensus box position 72), conserved as Tyr among mammalian Sry- and Sox domains (54). The first is Y127C. This *de novo* mutation presumably impairs specific DNA-dependent folding of the minor wing through three potential mechanisms: loss of side-chain volume leading to a destabilizing cavity (55), loss of favorable aromatic-aromatic interactions (56) and/or steric class at the side chain’s γ-position due to the larger atomic radius of sulfur versus carbon (57) (Fig. 6*B-D*). Y1H reporter assays suggested a marked loss of specific DNA-binding activity. In embryonic rat pre-Sertoli cells Y127C SRY underwent accelerated proteasomal degradation. Restoration of native protein expression by proteasome inhibitor MG132 did not rescue transcriptional activity (as monitored via autosomal target gene *Sox9*).

Y127C SRY is both abnormally susceptible to proteasomal degradation and intrinsically defective to transcriptional activity. Together with the Y1H results, these observations suggest that Y127C-associated impairment of specific DNA-dependent folding of the variant minor wing blocks SRY’s functional engagement at testis-specific target sites. It would be of future interest to survey the specific enhancer-binding properties (in cells) and specific DNA-binding properties (*in vitro*) of variant SRYs containing a diverse collection of substitutions at position 127. Our Y1H assay suggested that Trp127 would exhibit substantial activity (in accordance with occurrence of Trp at this site in SOX30; Fig. S2 (58)) but that Ala and His (not observed among mammalian Sry sequences or vertebrate Sox sequences (59)) would be almost as deleterious as Cys (Supplemental Fig. S2).

We chose to investigate in detail the biophysical properties and cellular function of inherited variant Y127F because of (a) its chemical subtlety and (b) a seeming paradox: whereas its inheritance demonstrates compatibility in principle with male development (in the proband’s father), a previous study reported no detectable specific DNA-binding activity (31). Such a developmental outcome suggests either (i) that the father exhibited WT germ-line mosaicism; (ii) that the SRY defect could be compensated or bypassed depending on genetic background or (iii) that the variant SRY gains a non-DNA-mediated function to effect male gene regulation. Although cryptic mosaicism cannot be excluded, the latter two suppositions seemed likely. Because the original report relied on the gel mobility-shift assay (GMSA)—susceptible to kinetic artifacts—we hypothesized that kinetic instability of the variant protein-DNA complex, if present, could have attenuated the shifted band as a known limitation of GMSA (23). In brief, because GMSA is not an equilibrium method, an accelerated rate of dissociation can lead to a diffuse or undetectable shifted band. In such cases GMSA data can underestimate the degree of residual DNA affinity (60).

Our biochemical and biophysical reassessment was further motivated by the substantial biological activity of Y127F SRY in an embryonic rat pre-Sertoli cell line (48). Indeed, under standard transfection conditions its transcriptional activation of endogenous target gene *Sox9* was indistinguishable from that of WT SRY (“1X” conditions in Fig. 2*A*). Reduction in expression levels into the physiological range uncovered a near-twofold attenuation of *Sox9* activation (“50X” conditions) in accordance with previous studies of diverse inherited Swyer alleles (28,33,34). This twofold transcriptional threshold is independent of the molecular or cellular mechanism of impairment (61). The substantial potency of Y127F SRY (consistent with that of other inherited alleles) indicates that cryptic mosaicism in the germ line need not be invoked to rationalize the phenotype of the proband’s father.

Equilibrium FRET-based measurement of specific DNA affinities demonstrated that Y72F impairs binding by only threefold at 37 °C. Stopped-flow kinetic studies nonetheless uncovered a sevenfold acceleration in *off* rate relative to dissociation of the WT complex. Together, these findings indicate that Y72F enhances the rate of binding by 2.5-fold (*k*_on_), which provides a partial kinetic compensation for the accelerated *off* rate (*k*_off_) to mitigate the net change in affinity (K_D_). DNA bend angles was provided by time-resolved FRET studies: the 15-bp DNA probe contained a fluorescent donor at one 5’-end (fluorescein) and an acceptor at the other 5’-end (TAMRA) as previously described (23). Whereas the variant’s mean reduction in end-to-end DNA distance was similar to that in the WT complex, the width of distances was subtly increased (Fig. 5*H*). It would be of future interest to obtain high-resolution crystal structures of the WT and variant domain-DNA complexes to gain insight into the structural origins of such tr-FRET findings.

We imagine that the *para*-OH group of Tyr127 functions in some way to enhance specific DNA binding and in particular to “lock” the complex. This lock has a kinetic aspect (retarding protein-DNA dissociation) and a dynamic aspect (enhancing the precision of DNA bending). The long-range kinetic and dynamic impact of a single functional group seems remarkable: how might this arise? We speculate that this Y127 *para*-OH group serves as an *anchor point* for a bound water molecule that bridges the protein and DNA surfaces. Although bound water molecules were not sought or identified in the NMR structure of the SRY box-DNA complex (21,22), an homologous protein-DNA co-crystal structure (containing the Sox2 HMG box) exhibits exactly such a bridging water molecule (Fig. 7; (46)). The minor wing and mini-core of the Sox2 HMG box are otherwise similar to those of the SRY domain. A corresponding water-mediated hydrogen bond also appears in the SOX18 co-crystal structure (62).

**Figure 7.**
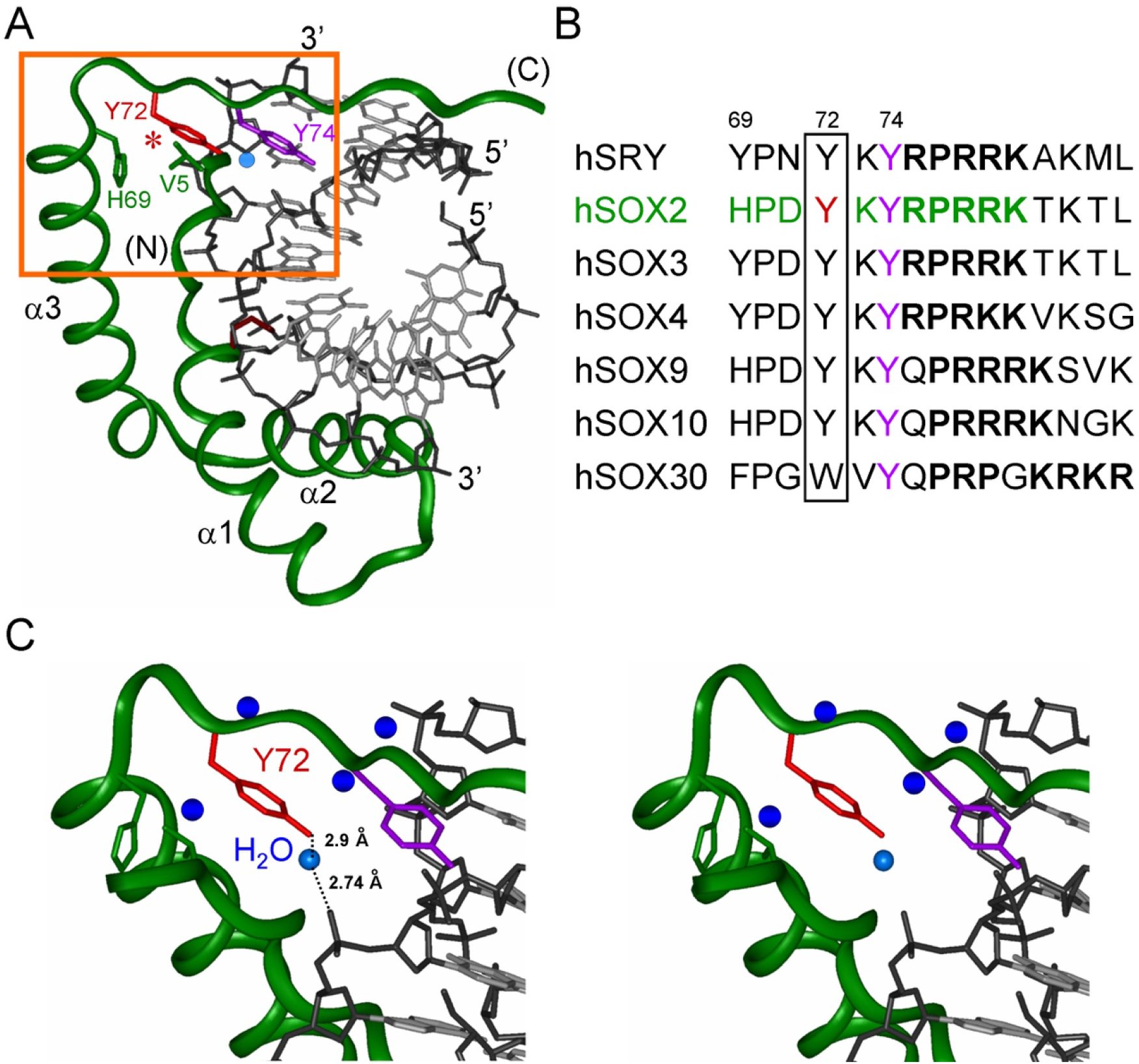
Tyr-anchored water-mediated hydrogen bond in homologous SOX2 complex. (*A*) Ribbon model of SOX2 HMG box; the domain docks within minor groove of bent DNA site (stick representation). Highlighted side chains in the ribbon are sites of minor wing (following residue numbers in consensus human SRY HMG box): *red*, Y72; and *green*, the remainder of the wing except the Y74 (*purple*). The DNA atoms are dark grey (phosphodiester linkages) and light gray (base pairs). Sphere highlighted with light blue is a bridging water molecule within the protein-DNA interface. The boxed region contains minor-wing-DNA contacts. (*B*) Sequence conservation of the minor wing of the human SRY/SOX HMG box. Tyr at position 72 (*boxed*) of the HMG box is highly conserved except SOX30. Side chain of Y74 (*purple*) of the minor wing is also conserved. Residues in position 69 are aromatic but mildly various. Residue consensus numbers following human SRY HMG box are provided above sequences. (*C*) Expansion of boxed region in panel *A* showing side-chain packing within the minor wing and protein-DNA contacts mediating by specific water molecule (oxygen molecule; *light blue*). Distances between DNA phosphodiester-water-hydroxyl group of Y72 are shown. Other oxygen of nearby water molecules are indicated with dark blue.

### Concluding remarks

Tyr and Phe differ only by the presence or absence of a *para*-OH group in corresponding aromatic rings. Our results demonstrate that the robustness of a human genetic switch can hinge on the presence of this hydroxyl group, which is solvent-exposed but not in direct contact with DNA. We propose that Tyr127 in human SRY (box position 72) has a dual function: first, to seal and stabilize the mini-core of the minor wing of the HMG box on specific DNA binding. Next, induced fit of the minor wing positions its *para*-OH at a favorable distance and orientation to enable a bound water molecule to contact a non-bridging oxygen atom in a phosphodiester DNA backbone. We envision that this bound water molecule, likely part of a solvation network at this interface, locks the bent protein-DNA complex to prolong its kinetic lifetime, augment its affinity and enhance its precision of DNA bending. Together, these favorable biophysical contributions enable WT SRY to cross the transcriptional threshold of testis determination. As foreshadowed by studies of the prokaryotic Trp repressor more than thirty years ago (63), bound water molecules can be critical elements in protein-DNA recognition. That loss of a single Tyr-anchored water molecule from the SRY-DNA interface can render human sex determination ambiguous seems remarkable.

## EXPERIMENTAL PROCEDURES

### Plasmids

Mammalian vector pCMX with the CMV promoter expressing full-length SRY variants containing a triplicate hemagglutinin (HA) tag were constructed as previously described (28). Bacterial plasmids expressing human SRY HMG boxes were constructed as described (28). For yeast-1-hybrid system, DNA encoding the human SRY HMG box was subcloned into plasmid pGAD-T7 to express fusion proteins following an N-terminal nuclear localization signal (NLS), central activation domain (AD) derived from GAL4, and C-terminal SRY fragment (28). Mutations in SRY were introduced using QuikChange™ (Stratagene). Constructions were verified by DNA sequencing.

### Y1H screening

Oligonucleotides containing the SRY triplicate consensus binding sites within the plasmid pLacZi (bold; 5’-AATTCGCA**ATTGTTATTGTTATTGTT**-3’) were constructed as described (28), and this site lies upstream of lacZ encoding β-galactosidase. Reporter strains bearing integrated SRY target or non-target sites were tested as described (28). Colonies were grown in minimal medium under selection. Extent of SRY-dependent expression of β-galactosidase was evaluated by quantitative assay of enzyme activity in liquid culture using *ortho*-nitrophenyl-β-galactoside (ONPG) as described by the vendor (Clontech Laboratories, Inc). *Mammalian Cell Culture—*Rodent preSertoli-cell line, CH34, kindly provided by T. R. Clarke and P. K. Donahoe (Massachusetts General Hospital, Boston, MA (20)), was employed for studies of SRY-regulation gene regulatory network. Cells were cultured in Dulbecco’s modified Eagle medium containing 5% heat-inactivated fetal bovine serum at 37 °C under 5% CO_2_. In studies using proteasome-inhibitor, transfected cells were maintained for 24 h in serum-free conditions and then treated with MG132 for 6 h followed by 18 h incubation in 5% serum-containing medium.

### Transient Transfection

Transfections were performed using Fugene HD (Hoffmann LaRoche) with protocols from vendor. Transfection efficiencies were determined by the co-transfection with pCMX-SRY and pCMX-GFP in equal amounts. Expression of the epitope-tagged SRY was monitored by Western blot. The epitope tag was provided by hemagglutinin (HA).

### Cycloheximide-chase assay

Post transfection cells were split evenly into 6-well plates and treated with translation inhibitor cycloheximide (final concentration is 20 mg/mL) in the regular medium for indicated times; cells were then lysed using mammalian lysis buffer (Hoffmann LaRoche). After normalization of total protein concentration, lysates were subjected to 12% SDS-PAGE and Western blot using anti-HA antiserum (Sigma-Aldrich) at a dilution ratio of 1:5000 with α-tubulin as a loading control. Quantification was performed by Image J program.

### Sox9 activation and downstream gene regulatory network

SRY-mediated transcriptional activation of Sox9 and other endogenous CH34 genes was measured in triplicate by quantitative real-time-quantitative-rtPCR (qPCR) as described (28). Cellular RNA was extracted using RNeasy (Qiagen).

### Immunocytochemistry

Transfected cells were plated evenly on 12-mm cover slips, fixed with 3% *para*-formaldehyde in phosphate-buffered saline (PBS) and then treated with permeability buffer solution (PBS containing 10% goat serum and 1% Triton X-100; Sigma-Aldrich). SRY variants were probed using FITC-conjugated anti-HA antibody (Santa Cruz). cells were visualized by fluorescent microscopy; DAPI stained the nucleus. Nuclear localization was evaluated by the ratio of cells expressing SRY in the nucleus to the total number of GFP-positive cells.

### Chromatin immunoprecipitation

Cells were transfected with SRY variants and cross-linked in wells by formaldehyde, collected, and lysed after quenching the cross-linking reaction. Chromatin lysates were sonicated for generating 300-400-bp fragments which were immunoprecipitated using anti-HA antiserum (Sigma-Aldrich) coupled with Protein A slurry (Santa Cruz). Non-specific antiserum (Santa Cruz) served as control. After reversal of cross-linking, fragments were treated with proteinase K and RNase (Hoffmann LaRoche), followed by extraction using 1:1 phenol-CIAA solution. PCR protocol was provided by the vender (Hoffmann LaRoche).

### Fluorescence and Circular Dichroism spectroscopy

Stability of the free domain was determined using fluorescence spectroscopy and monitoring the Trp emission wavelength 390nm. Free protein domains were made 5uM in Buffer A (10mM potassium phosphate buffer pH 7.4 and 140 mM KCl). For guanidine titration, 5 μM protein was made with same buffer components but dissolved in 8 M guanidine hydrochloride. Protein-DNA complex melting temperature (T_m_) were measured by CD at α-helical wavelength 222 nm. Protein-DNA complexes were made in Buffer A at 25 μM and data collected from 4-95 °C.

### FRET based assays

A 15-base pair (bp) DNA duplex was fluorescently modified at the 5’-end of each strand to contain either: fluorescein (donor) or tetramethylrhodamine (TAMRA; acceptor). Kinetic off rates (*k*_*off*_) were determined using stopped-flow FRET wherein a solution containing an equimolar labeled-DNA and protein was rapidly mixed with a solution containing an unmodified 15-bp DNA target. Donor emission was monitored at 520 nm, and *off*-rate constants determined using a single-exponential decay model. Specific DNA binding (K_D_) was determined using a constant concentration of modified DNA (25 nM) with increasing concentrations of the proteins. The change in the FRET signals with increasing protein concentration were plotted and fit as described (23).

### Time-resolved FRET assays

In house-built system at Bar Ilan University employs time-correlated single-photon counting as described (64). Emission wavelength was selected by a double 0.125m subtractive monochromator. The donor emission was collected at 520 nm. Distance distributions were obtained from global analysis of experimental decay curves using Marquardt non-linear least squares (65). Measurements were conducted in triplicate.

## Abbreviations

Bp: base pair
CD: circular dichroism
ChIP: chromatin immunoprecipitation
FRET: fluorescence-resonance energy transfer
GMSA: gel mobility-shift assay
GFP: green fluorescent protein
GuHC: guanidine hydrochloride
HA: hemagglutinin
PCR: polymerase chain reaction
TAMRA: tetramethylrhodamine
tr-FRET: time-resolved FRET
WT: wild-type
Y1H: yeast-one-hybrid. Amino acids are designated by standard one- and three-letter codes

## Contributions

Cell-based assays were performed by Y.-S.C. Steady-state FRET studies and kinetic analyses were performed by J.R. Time-resolved FRET studies were performed and analyzed by D.A. and E.H. The overall study was coordinated by M.A.W.

## Acknowledgements

We thank Drs. N. Phillips (Case Western Reserve University School of Medicine) and M. Georgiadis (Indiana University School of Medicine) for helpful discussions. This work was funded in part by a grant to MAW from the National Institutes of Health.

## Notes

### Competing Interest Statement

The authors have declared no competing interest.

## REFERENCES

1. Sinclair, A. H., Berta, P., Palmer, M. S., Hawkins, J. R., Griffiths, B. L., Smith, M. J., Foster, J. W., Frischauf, A. M., Lovell-Badge, R., and Goodfellow, P. N. (1990) A gene from the human sex-determining region encodes a protein with homology to a conserved DNA-binding motif. Nature 346, 240–244

2. Sekido, R. (2010) SRY: a transcriptional activator of mammalian testis determination. Int. J. Biochem. Cell Biol. 42, 417–420

3. Ner, S. S. (1992) HMGs everywhere. Curr. Biol. 2, 208–210

4. Berta, P., Hawkins, J. R., Sinclair, A. H., Taylor, A., Griffiths, B. L., Goodfellow, P. N., and Fellous, M. (1990) Genetic evidence equating SRY and the testis-determining factor. Nature 348, 448–450

5. Knower, K. C., Kelly, S., and Harley, V. R. (2003) Turning on the male--SRY, SOX9 and sex determination in mammals. Cytogenet. Genome Res. 101, 185–198

6. Hawkins, J. R., Taylor, A., Berta, P., Levilliers, J., Van der Auwera, B., and Goodfellow, P. N. (1992) Mutational analysis of SRY: nonsense and missense mutations in XY sex reversal. Hum. Genet. 88, 471–474

7. Cameron, F. J., and Sinclair, A. H. (1997) Mutations in SRY and SOX9: testis-determining genes. Hum. Mutat. 9, 388–395

8. Koopman, P., Gubbay, J., Vivian, N., Goodfellow, P., and Lovell-Badge, R. (1991) Male development of chromosomally female mice transgenic for Sry. Nature 351, 117–121

9. Sekido, R., Bar, I., Narvaez, V., Penny, G., and Lovell-Badge, R. (2004) SOX9 is up-regulated by the transient expression of SRY specifically in Sertoli cell precursors. Dev. Biol. 274, 271–279

10. Sekido, R., and Lovell-Badge, R. (2008) Sex determination involves synergistic action of SRY and SF1 on a specific Sox9 enhancer. Nature 453, 930–934

11. Gonen, N., Futtner, C. R., Wood, S., Garcia-Moreno, S. A., Salamone, I. M., Samson, S. C., Sekido, R., Poulat, F., Maatouk, D. M., and Lovell-Badge, R. (2018) Sex reversal following deletion of a single distal enhancer of Sox9. Science 360, 1469–1473

12. Kobayashi, A., Chang, H. A. O., Chaboissier, M. C., Schedl, A., and Behringer, R. R. (2005) Sox9 in testis determination. Ann. N. Y. Acad. Sci. 1061, 9–17

13. Lovell-Badge, R., Canning, C., and Sekido, R. (2002) Sex-determining genes in mice: building pathways, John Wiley & Sons Ltd., West Sussex, UK

14. Josso, N., Picard, J. Y., Rey, R., and di Clemente, N. (2006) Testicular anti-Mullerian hormone: history, genetics, regulation and clinical applications. Pediatr. Endocrinol. Rev. 3, 347–358

15. MacLaughlin, D. T., and Donahoe, P. K. (2004) Sex determination and differentiation. N. Engl. J. Med. 350, 367–378

16. McClelland, K., Bowles, J., and Koopman, P. (2012) Male sex determination: insights into molecular mechanisms. Asian J. Androl. 14, 164–171

17. Wilhelm, D., and Koopman, P. (2006) The makings of maleness: towards an integrated view of male sexual development. Nat. Rev. Genet. 7, 620–631

18. Anu, B., and Ken, M. (2015) Human sex-determination and disorders of sex-development (DSD). Seminars in Cell & Developmental Biology 45, 77–83

19. Pontiggia, A., Rimini, R., Harley, V. R., Goodfellow, P. N., Lovell-Badge, R., and Bianchi, M. E. (1994) Sex-reversing mutations affect the architecture of SRY-DNA complexes. EMBO J. 13, 6115–6124

20. Haqq, C. M., King, C. Y., Ukiyama, E., Falsafi, S., Haqq, T. N., Donahoe, P. K., and Weiss, M. A. (1994) Molecular basis of mammalian sexual determination: activation of Müllerian inhibiting substance gene expression by SRY. Science 266, 1494–1500

21. Werner, M. H., Bianchi, M. E., Gronenborn, A. M., and Clore, G. M. (1995) NMR spectroscopic analysis of the DNA conformation induced by the human testis determining factor SRY. Biochemistry 34, 11998–12004

22. Murphy, E. C., Zhurkin, V. B., Louis, J. M., Cornilescu, G., and Clore, G. M. (2001) Structural basis for SRY-dependent 46-X,Y sex reversal: modulation of DNA bending by a naturally occuring point mutation. J. Mol. Biol. 312, 481–499

23. Phillips, N. B., Jancso-Radek, A., Ittah, V., Singh, R., Chan, G., Haas, E., and Weiss, M. A. (2006) SRY and human sex determination: The basic tail of the HMG box functions as a kinetic clamp to augment DNA bending. J. Mol. Biol. 358, 172–192

24. Bewley, C. A., Gronenborn, A. M., and Clore, G. M. (1998) Minor groove-binding architectural proteins: structure, function, and DNA recognition. Annu. Rev. Biophys. Biomol. Struct. 27, 105–131

25. Weiss, M. A. (2005) Molecular mechanisms of male sex determination: the enigma of SRY. in DNA Conformation and Transcription (Ohyama, T. ed.), Landes Bioscience, Georgetown, TX. pp 1–15

26. Sekido, R., and Lovell-Badge, R. (2009) Sex determination and SRY: down to a wink and a nudge? Trends Genet. 25, 19–29

27. Kashimada, K., and Koopman, P. (2010) SRY: the master switch in mammalian sex determination. Development 137, 3921–3930

28. Phillips, N. B., Racca, J., Chen, Y. S., Singh, R., Jancso-Radek, A., Radek, J. T., Wickramasinghe, N. P., Haas, E., and Weiss, M. A. (2011) Mammilian testis-determining factor SRY and the enigma of inherited human sex reversal. J. Biol. Chem. 286, 36787–36807

29. Harley, V. R., Jackson, D. I., Hextall, P. J., Hawkins, J. R., Berkovitz, G. D., Sockanathan, S., Lovell-Badge, R., and Goodfellow, P. N. (1992) DNA binding activity of recombinant SRY from normal males and XY females. Science 255, 453–456

30. Dork, T., Stuhrmann, M., Miller, K., and Schmidtke, J. (1998) Independent observation of SRY mutation I90M in a patient with complete gonadal dysgenesis. Hum. Mutat. 11, 90–91

31. Jordan, B. K., Jain, M., Natarajan, S., Frasier, S. D., and Vilain, E. (2002) Familial mutation in the testis-determining gene SRY shared by an XY female and her normal father. J. Clin. Endocrinol. Metab. 87, 3428–3432

32. Domenice, S., Yumie Nishi, M., Correia Billerbeck, A. E., Latronico, A. C., Aparecida Medeiros, M., Russell, A. J., Vass, K., Marino Carvalho, F., Costa Frade, E. M., Prado Arnhold, I. J., and Bilharinho Mendonca, B. (1998) A novel missense mutation (S18N) in the 5’ non-HMG box region of the SRY gene in a patient with partial gonadal dysgenesis and his normal male relatives. Hum. Genet. 102, 213–215

33. Chen, Y. S., Racca, J. D., Phillips, N. B., and Weiss, M. A. (2013) Inherited human sex reversal due to impaired nucleocytoplasmic trafficking of SRY defines a male transcriptional threshold. Proc. Natl. Acad. Sci. U. S. A. 110, E3567–3576

34. Racca, J. D., Chen, Y.-S., Yang, Y., Phillips, N. B., and Weiss, M. A. (2016) Human sex determination at the edge of ambiguity inherited XY sex reversal due to enhanced ubiquitination and proteasomal degradation of a master transcription factor. Journal of Biological Chemistry 291, 22173–22195

35. Weir, H. M., Kraulis, P. J., Hill, C. S., Raine, A. R., Laue, E. D., and Thomas, J. O. (1993) Structure of the HMG box motif in the B-domain of HMG1. EMBO J. 12, 311–319

36. Read, C. M., Cary, P. D., Crane-Robinson, C., Driscoll, P. C., and Norman, D. G. (1993) Solution structure of a DNA-binding domain from HMG1. Nucleic Acids Res. 21, 3427–3436

37. Jones, D. N., Searles, M. A., Shaw, G. L., Churchill, M. E., Ner, S. S., Keeler, J., Travers, A. A., and Neuhaus, D. (1994) The solution structure and dynamics of the DNA-binding domain of HMG-D from Drosophila melanogaster. Structure 2, 609–627

38. van Houte, L. P., Chuprina, V. P., van der Wetering, M., Boelens, R., Kaptein, R., and Clevers, H. (1995) Solution structure of the sequence-specific HMG box of the lymphocyte transcriptional activator Sox-4. J. Biol. Chem. 270, 30516–30524

39. Weiss, M. A. (2001) Floppy SOX: mutual induced fit in hmg (high-mobility group) box-DNA recognition. Mol. Endocrinol. 15, 353–362

40. Li, B., Phillips, N. B., Jancso-Radek, A., Ittah, V., Singh, R., Jones, D. N., Haas, E., and Weiss, M. A. (2006) SRY-directed DNA bending and human sex reversal: reassessment of a clinical mutation uncovers a global coupling between the HMG box and its tail. J. Mol. Biol. 360, 310–328

41. Werner, H. M., Huth, J. R., Gronenborn, A. M., and Clore, G. M. (1995) Molecular basis of human 46X,Y sex reversal revealed from the three-dimensional solution structure of the human SRY-DNA complex. Cell 81, 705–714

42. Laudet, V., Stehelin, D., and Clevers, H. (1993) Ancestry and diversity of the HMG box superfamily. Nucleic Acids Res. 21, 2493–2501

43. Braun. (1993) True hermaphroditism in a 46,XY individual, caused by a postzygotic somatic point mutation in the male gonadal sex-determining locus (SRY): molecular genetics and histological findings in a sporadic case. Am. J. Hum. Genet. 52, 578–585

44. Hiort, O., Gramss, B., and Klauber, G. T. (1995) True hermaphroditism with 46, XY karyotype and a point mutation in the SRY gene. J. Pediatr. 126, 1022

45. Poulat, F., Soullier, S., Goze, C., Heitz, F., Calas, B., and Berta, P. (1994) Description and functional implications of a novel mutation in the sex-determining gene SRY. Hum. Mutat. 3, 200–204

46. Reményi, A., Lins, K., Nissen, L. J., Reinbold, R., Schöler, H. R., and Wilmanns, M. (2003) Crystal structure of a POU/HMG/DNA ternary complex suggests differential assembly of Oct4 and Sox2 on two enhancers. Genes & development 17, 2048–2059

47. Klaus, M., Prokoph, N., Girbig, M., Wang, X., Huang, Y.-H., Srivastava, Y., Hou, L., Narasimhan, K., Kolatkar, P. R., and Francois, M. (2016) Structure and decoy-mediated inhibition of the SOX18/Prox1-DNA interaction. Nucleic acids research, gkw130

48. Haqq, C. M., and Donahoe, P. K. (1998) Regulation of sexual dimorphism in mammals. Physiol. Rev. 78, 1–33

49. Koopman, P., Bullejos, M., and Bowles, J. (2001) Regulation of male sexual development by Sry and Sox9. J. Exp. Zool. 290, 463–474

50. Weiss, M. A., Ukiyama, E., and King, C. Y. (1997) The SRY cantilever motif discriminates between sequence-and structure-specific DNA recognition: alanine mutagenesis of an HMG box. J. Biomol. Struct. Dyn. 15, 177–184

51. Bullejos, M., and Koopman, P. (2005) Delayed Sry and Sox9 expression in developing mouse gonads underlies B6-YDOM sex reversal. Dev. Biol. 278, 473–481

52. Moniot, B., Declosmenil, F., Barrionuevo, F., Scherer, G., Aritake, K., Malki, S., Marzi, L., Cohen-Solal, A., Georg, I., Klattig, J., Englert, C., Kim, Y., Capel, B., Eguchi, N., Urade, Y., Boizet-Bonhoure, B., and Poulat, F. (2009) The PGD2 pathway, independently of FGF9, amplifies SOX9 activity in Sertoli cells during male sexual differentiation. Development 136, 1813–1821

53. Schmitt-Ney, M., Thiele, H., Kaltwasser, P., Bardoni, B., Cisternino, M., and Scherer, G. (1995) Two novel SRY missense mutations reducing DNA binding identified in XY females and their mosaic fathers. Am. J. Hum. Genet. 56, 862–869

54. Pontiggia, A., Whitfield, S., Goodfellow, P. N., Lovell-Badge, R., and Bianchi, M. E. (1995) Evolutionary conservation in the DNA-binding and -bending properties of HMG-boxes from SRY proteins of primates. Gene 154, 277–280

55. Racca, J. D., Chen, Y.-S., Maloy, J. D., Wickramasinghe, N., Phillips, N. B., and Weiss, M. A. (2014) Structure-Function Relationships in Human Testis-determining Factor SRY: AN AROMATIC BUTTRESS UNDERLIES THE SPECIFIC DNA-BENDING SURFACE OF A HIGH MOBILITY GROUP (HMG) BOX. Journal of Biological Chemistry 289, 32410–32429

56. Serrano, L., Bycroft, M., and Fersht, A. R. (1991) Aromatic-aromatic interactions and protein stability: investigation by double-mutant cycles. Journal of molecular biology 218, 465–475

57. Grutter, M. G., Gray, T. M., Weaver, L. H., Wilson, T. A., and Matthews, B. W. (1987) Structural studies of mutants of the lysozyme of bacteriophage T4. The temperature-sensitive mutant protein Thr157 le. J. Mol. Biol. 197, 315–329

58. Han, F., Wang, Z., Wu, F., Liu, Z., Huang, B., and Wang, D. (2010) Characterization, phylogeny, alternative splicing and expression of Sox30 gene. BMC Molecular Biology 11, 98

59. Bowles, J., Schepers, G., and Koopman, P. (2000) Phylogeny of the SOX family of developmental transcription factors based on sequence and structural indicators. Dev. Biol. 227, 239–255

60. Ross, J. R., Martha, P. B., Adeola, G., Deanna, R., and Catherine, A. R. (1997) Affinity and specificity of trp repressor-DNA interactions studied with fluorescent oligonucleotides11Edited by P.E. Wright. Journal of Molecular Biology 273, 572–585

61. Chen, Y.-S., Racca, J. D., Sequeira, P. W., and Weiss, M. A. (2015) Inherited Sex-Reversal Mutations in SRY Define a Functional Threshold of Gonadogenesis: Biochemical and Evolutionary Implications of a Rare Monogenic Syndrome. Rare Disorders: Diagnosis & Therapy

62. Klaus, M., Prokoph, N., Girbig, M., Wang, X., Huang, Y. H., Srivastava, Y., Hou, L., Narasimhan, K., Kolatkar, P. R., Francois, M., and Jauch, R. (2016) Structure and decoy-mediated inhibition of the SOX18/Prox1-DNA interaction. Nucleic Acids Res 44, 3922–3935

63. Otwinowski, Z., Schevitz, R. W., Zhang, R. G., Lawson, C. L., Joachimiak, A., Marmorstein, R. Q., Luisi, B. F., and Sigler, P. B. (1988) Crystal structure of trp repressor/operator complex at atomic resolution. Nature 335, 321–329

64. Phillips, N. B., Nikolskaya, T., Jancso-Radek, A., Ittah, V., Jiang, F., Singh, R., Haas, E., and Weiss, M. A. (2004) Sry-directed sex reversal in transgenic mice is robust to enhanced DNA bending: comparison of human and murine in HMG boxes. Biochemistry 43, 7066–7081

65. Beechem, J. M., and Haas, E. (1989) Simultaneous determination of intramolecular distance distributions and conformational dynamics by global analysis of energy transfer measurements. Biophys. J. 55, 1225–1236

66. Desclozeaux, M., Poulat, F., de Santa Barbara, P., Capony, J.-P., Turowski, P., Jay, P., Méjean, C., Moniot, B., Boizet, B., and Berta, P. (1998) Phosphorylation of an N-terminal motif enhances DNA-binding activity of the human SRY protein. Journal of Biological Chemistry 273, 7988–7995

67. Poulat, F., de Santa Barbara, P., Desclozeaux, M., Soullier, S., Moniot, B., Bonneaud, N., Boizet, B., and Berta, P. (1997) The human testis determining factor SRY binds a nuclear factor containing PDZ protein interaction domains. Journal of Biological Chemistry 272, 7167–7172

68. King, C. Y., and Weiss, M. A. (1993) The SRY high-mobility-group box recognizes DNA by partial intercalation in the minor groove: a topological mechanism of sequence specificity. Proc. Natl. Acad. Sci. U. S. A. 90, 11990–11994

