## supplemental figures 1 and 2 for "Inherited Human Sex Reversal due to Loss of a Water-Mediated Hydrogen Bond at a Conserved Protein-DNA Interface"

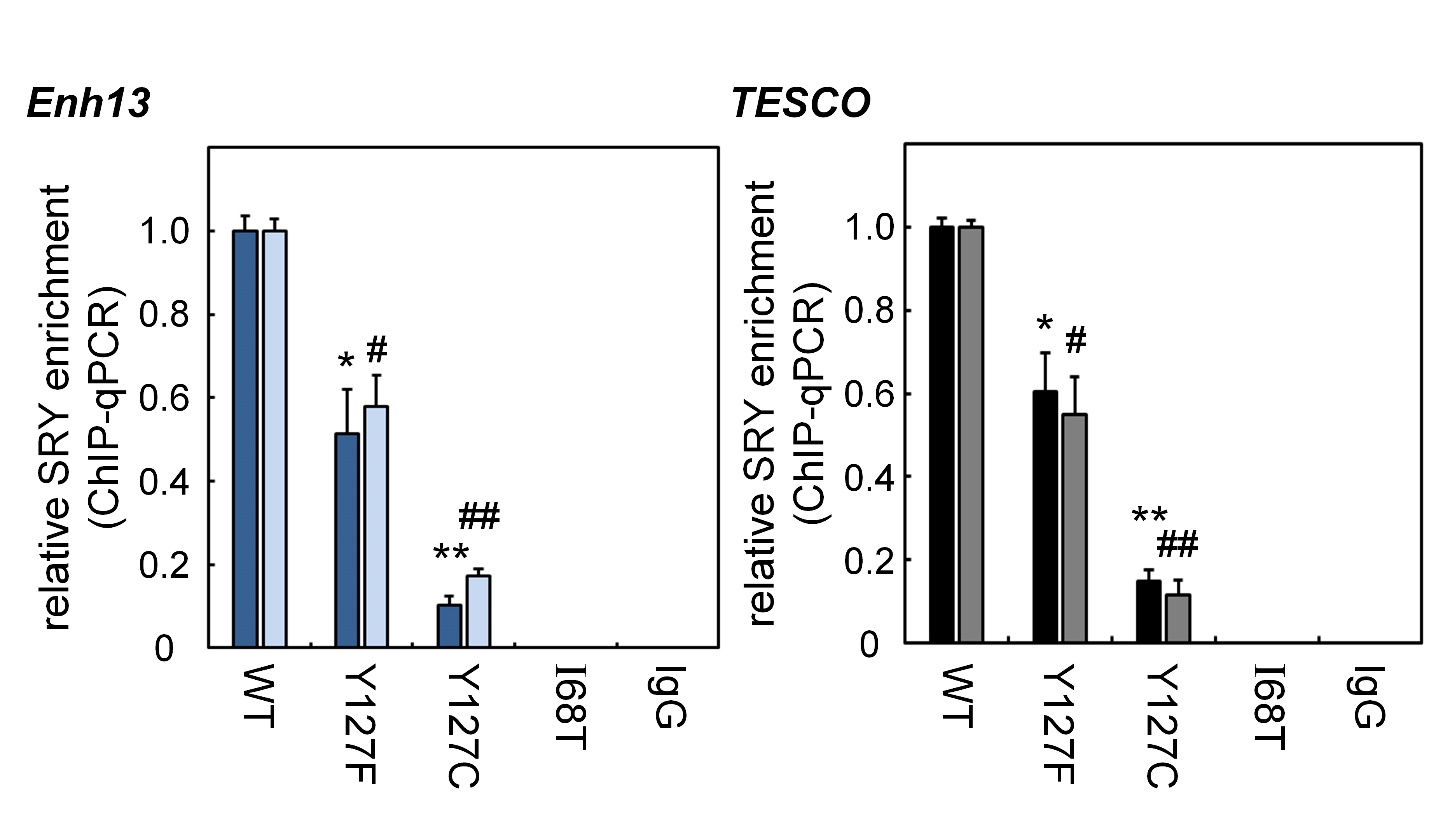

**Figure S1.** Site variants on SRY affect *Sox9* enhancer occupancy. *Sox9* gene is regulated by a far upstream enhancer element (*Enh13*) and *TES* core elements (*TESCO*) (11). Potential SRY binding sites are in *Enh13* fragments 1 and 3 and *TESCO* fragments 4 and 8. These fragments served as probes for monitoring SRY-enhancer occupancies. SRY-Enh13 (*Left*) and SRY-TESCO (*Right*) complexes were probed by anti-HA WB in ChIP assays quantified by qPCR; results are summarized by histogram. *Filled* and *open* bars represent occupancies of *Enh13* fragment 1 and 3, and *TESCO* fragment 4 and 8, respectively. Bar pairs correspond to WT SRY and Y127 variants. Control lane I68A and IgG indicate inactive I68A SRY and non-specific control. (*) and (#) indicate the statistical significance at the level p<0.05 (Wilcox test) for the occupancies.

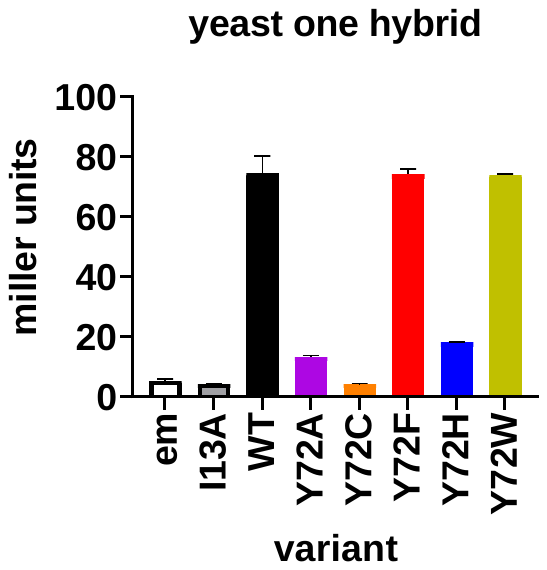

**Figure S2.** Yeast one hybrid results. Five Y72 variants were assayed for retained sequence-specific DNA binding in an SRY HMG-sensitive yeast reporter strain. Empty vector (em) and I13A serve as negative controls compared to wild type (WT).
